## Supplemental Information for "Analysis of the Immune Response to Sciatic Nerve Injury Identifies Efferocytosis as a Key Mechanism of Nerve Debridement"

Figure 1 – Figure Supplement 1

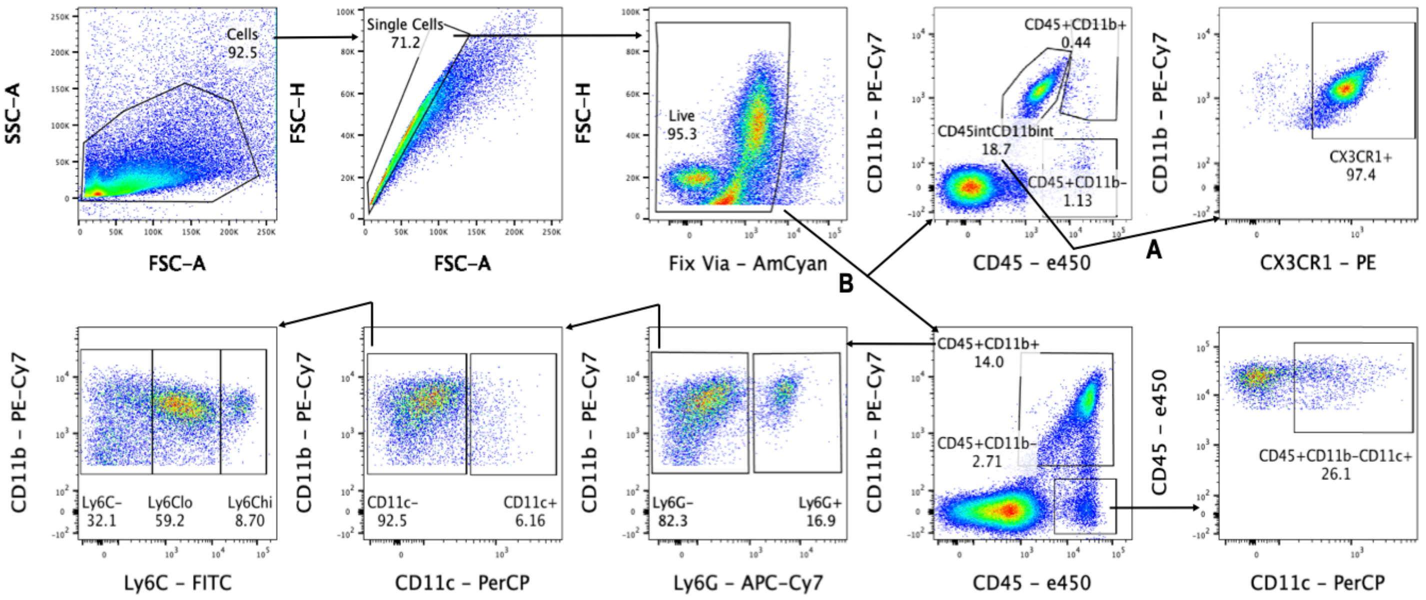

**Figure 1 – figure supplement 1. Gating scheme for flow cytometry.**

Cells were first gated with forward scatter (FSC-A) and side scatter (SSC-A) to exclude debris. Cells were then gated with forward scatter height (FSC-H) and FSC-A to find single cells and to exclude doublets. Live cells were isolated by negative staining for fixed viability dye (Fix Via). In nerves, DRGs, spinal cord, and spleen, leukocytes were analyzed as follows: lymphocytes were isolated as $CD45^+$, $CD11b^-$ cell and then further separated based on CD11c positivity as cDC or as $CD45^+$, $CD11b^-$, $CD11c^-$ lymphocytes. Myeloid cells ($CD45^+$, $CD11b^+$) were further separated into $Ly6G^+$ granulocytes. The remaining cells ($CD45^+$, $CD11b^+$, $Ly6G^-$) were characterized as MoDC ($CD45^+$, $CD11b^+$, $CD11c^+$, $Ly6G^-$), Mo/Mac ($CD45^+$, $CD11b^+$, $CD11c^-$, $Ly6G^-$), and Microglia ($CD45^{int}$, $CD11b^{int}$, $CX3CR1^+$).

Figure 1 – Figure Supplement 2

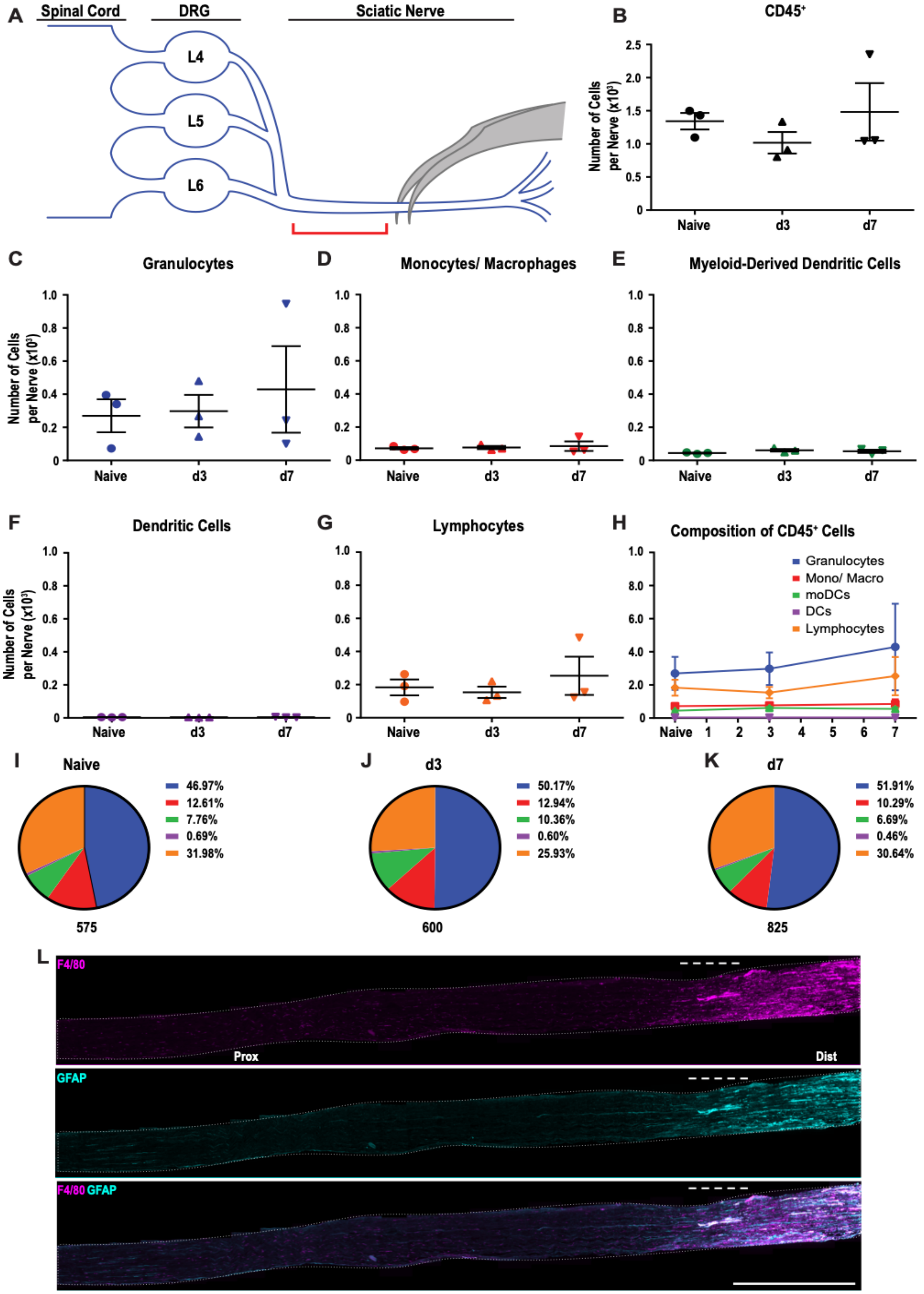

**Figure 1 – figure supplement 2. Immune cell profiles in the sciatic nerve proximal to the injury site.**

**A.** Anatomy of lumbar spinal cord and DRGs connected to the sciatic nerve. The location of the crush site within the nerve trunk and the tissue segment collected for flow cytometry (red bracket) are shown. **B**. Quantification of live, $CD45^+$ cell in the proximal nerve, per ~ 5mm segment. Flow cytometry of nerve tissue collected from naïve mice (n= 3), d3 (n= 3), and d7 (n= 3) following SNC. **C**. Quantification of granulocytes ($CD45^+$, $CD11b^+$, $Ly6G^+$) per nerve segment. **D**. Quantification of Mo/Mac ($CD45^+$, $CD11b^+$, $CD11c^-$, $Ly6G^-$) per nerve segment. **E**. Quantification of MoDC ($CD45^+$, $CD11b^+$, $CD11c^+$, $Ly6G^-$) per nerve segment **F.** Quantification of cDC ($CD45^+$, $CD11b^-$, $CD11c^+$, $Ly6G^-$) per nerve segment. **G**. Quantification of lymphocytes ($CD45^+$, $CD11b^-$) per nerve segment. **H**. Composition of $CD45^+$ leukocytes in the proximal nerve stump at different post-SNC time points. **I-K.** Percentile of each cell type at different post-injury time points. For flow cytometry, data are represented as mean cell number ± SEM. Statistical analysis was performed in GraphPad Prism (v7) using 1-way or 2-way ANOVA with correction for multiple comparisons with Tukey's post-hoc test. For B-G, unpaired two-tailed t-test with Welch's correction. A $p$ value $< 0.05$ (*) was considered significant. $p < 0.01$ (**), $p < 0.001$ (***), and $p < 0.0001$ (****). **L**. Longitudinal sections of sciatic nerve trunk at d3 following SNC, stained with anti-F4/80 (red) to label macrophages and anti-GFAP (green) to label repair Schwann cells. The proximal (Prox) and distal (Dist) sides of the nerve, relative to the crush site (dashed line), are indicated. Consistent with flow cytometry, nerve inflammation is not significantly elevated proximal to the nerve crush site. Scale bar, 1000 μm.

Figure 2 – Figure Supplement 1

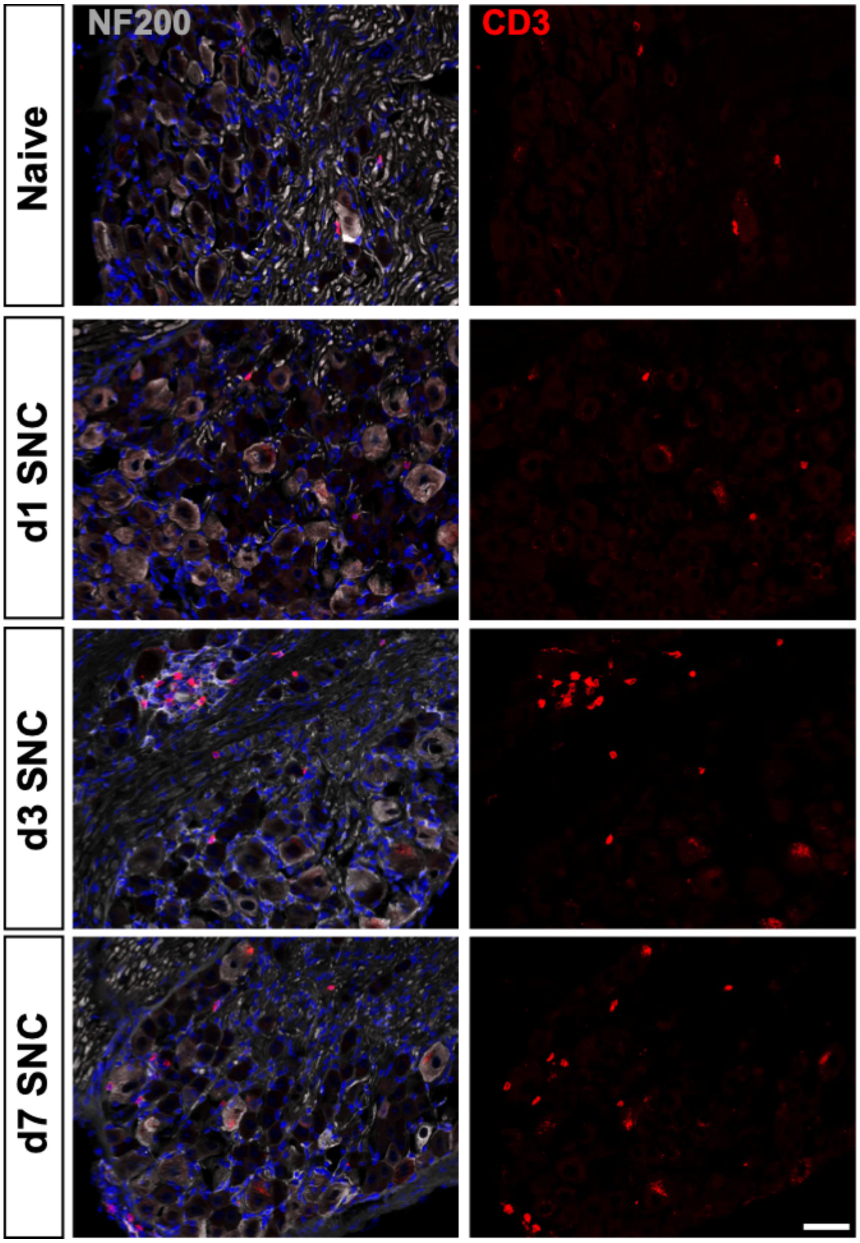

**Figure 2 – figure supplement 1. T cells in naïve and axotomized DRGs.**

Representative images of L4 DRG cross sections from naïve mice, d1, d3, and d7 post-SNC. T cells were labeled with anti-CD3 (red), neurons with anti-NF200 (white). DAPI (blue) was use for nuclear staining (blue). Scale bar, 50 μm.

Figure 3 – Figure Supplement 1

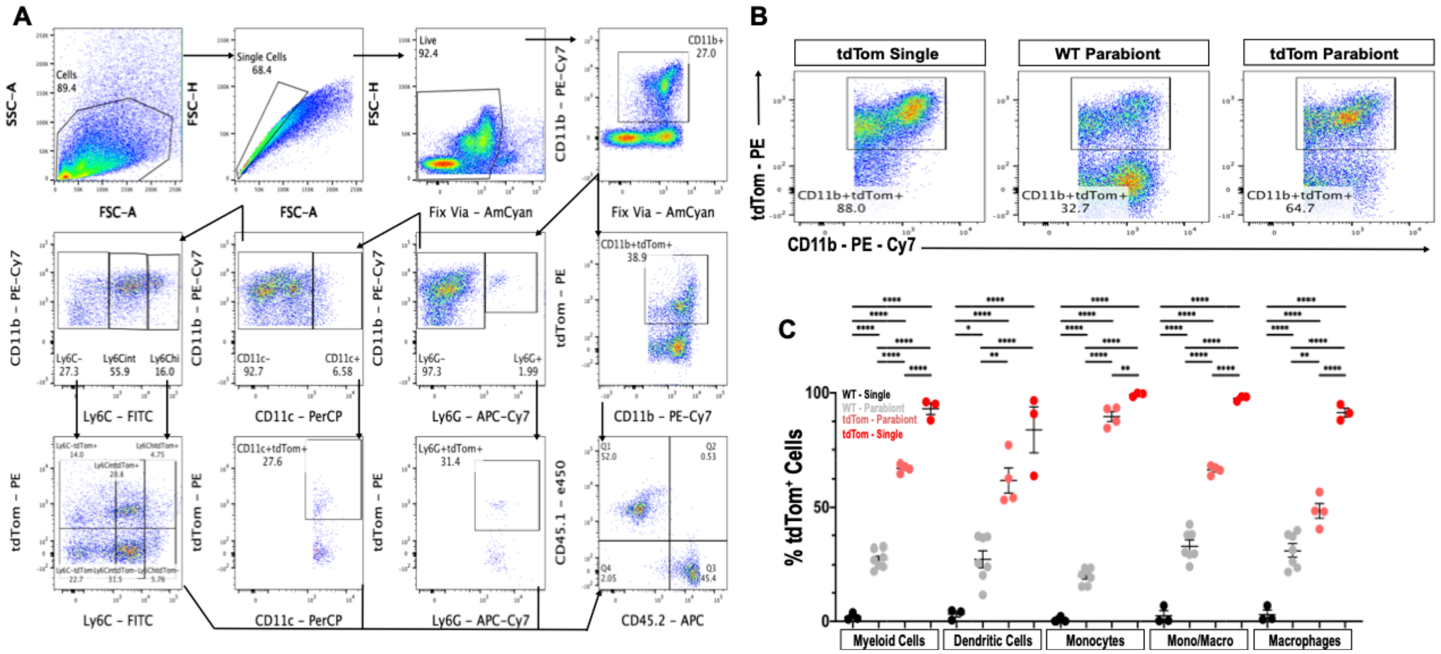

**Figure 3 – figure supplement 1. Flow cytometry gating scheme to assess chimerism of parabiotic mice.**

**A**. Cells were first gated with forward scatter (FSC-A) and side scatter (SSC-A) to exclude debris. Cells were then gated with forward scatter height (FSC-H) and FSC-A to find single cells and to exclude doublets. Live cells were isolated by negative staining for fixed viability dye (Fix Via). After live cell gating, myeloid cells were isolated by CD11b positivity and assessed for the percentage of $tdTom^+$ cells. Following gating for granulocytes ($CD11b^+$, $Ly6G^+$), MoDC ($CD11b^+$, $CD11c^+$, $Ly6G^-$), and Mo/Mac ($CD11b^+$, $Ly6G^-$, $CD11c^-$) were identified. Mo/Mac were further subdivided into $Ly6C^{hi}$, $Ly6C^{int}$ and $Ly6C^-$ cells. The fraction of $tdTom^+$ cells for each cell type was determined. **B**. Representative flow cytometry dot plots of splenic myeloid cells ($CD11b^+$, $tdTom^+$) in WT (CD45.1) parabiont and non-parabiotic (single) $tdTom^+$ and parabiotic mice. **C.** Quantification of chimerism in the spleen. The percentile of $tdTom^+$ myeloid cells, MoDC, Mo/Mac that are ($Ly6C^{hi}$), ($Ly6C^{int}$), and Mac ($Ly6C^-$) is shown (n= 3-7 biological replicas). Flow data are represented as mean ± SEM. Statistical analysis was performed in GraphPad Prism (v8) using 1-way ANOVA with correction for multiple comparisons with Tukey's post-hoc test. A p value $< 0.05$ (*) was considered significant. $p < 0.01$ (**), and $p < 0.0001$ (****).

Figure 4 – Figure Supplement 1

A

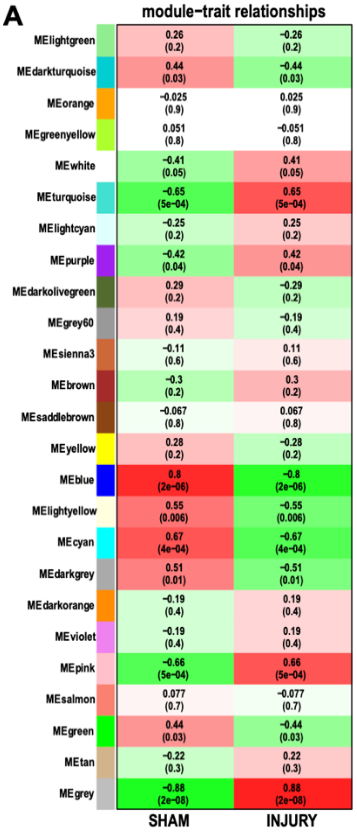


B

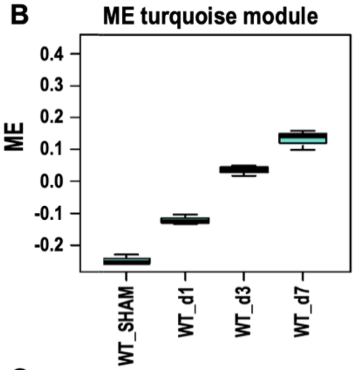


C

| X | NES |
|---|---|
| GO_CELL_ACTIVATION | 5.491423 |
| GO_IMMUNE_EFFECTOR_PROCESS | 5.456118 |
| GO_DEFENSE_RESPONSE | 5.291974 |
| GO_EXTRACELLULAR_STRUCTURE_ORGANIZATION | 5.268342 |
| GO_REGULATION_OF_IMMUNE_SYSTEM_PROCESS | 5.024953 |
| GO_INNATE_IMMUNE_RESPONSE | 4.932145 |
| GO_MYELOID_LEUKOCYTE_ACTIVATION | 4.905169 |
| GO_COLLAGEN_CONTAINING_EXTRACELLULAR_MATRIX | 4.768177 |
| GO_BIOLOGICAL_ADHESION | 4.733691 |
| GO_REGULATION_OF_CELL_POPULATION_PROLIFERATION | 4.60847 |
| GO_INTERSPECIES_INTERACTION_BETWEEN_ORGANISMS | 4.554828 |
| GO_EXTRACELLULAR_MATRIX | 4.534409 |
| GO_LEUKOCYTE_MEDIATED_IMMUNITY | 4.523114 |
| GO_CELL_ACTIVATION_INVOLVED_IN_IMMUNE_RESPONSE | 4.499422 |
| GO_RESPONSE_TO_CYTOKINE | 4.411848 |
| GO_PHAGOCYTOSIS | 4.382074 |
| GO_DNA_PACKAGING_COMPLEX | 4.380028 |
| GO_EXTRACELLULAR_MATRIX_STRUCTURAL_CONSTITUENT | 4.342254 |
| GO_LEUKOCYTE_MIGRATION | 4.239735 |

D

| Upstream Regulator | Molecule Type | p-value of overlap |
|---|---|---|
| lipopolysaccharide | chemical drug | 2.76E-145 |
| IFNG | cytokine | 2.32E-135 |
| TNF | cytokine | 1.82E-108 |
| IL1B | cytokine | 2.51E-86 |
| poly rI:rC-RNA | biologic drug | 8.88E-81 |
| STAT1 | transcription regulator | 1.43E-71 |
| STAT3 | transcription regulator | 1.21E-70 |
| resiquimod | chemical drug | 1.68E-70 |
| IL6 | cytokine | 5.14E-68 |
| TGFB1 | growth factor | 5.54E-65 |
| IL4 | cytokine | 3.46E-64 |
| dexamethasone | chemical drug | 5.03E-63 |
| NFkB (complex) | complex | 4.70E-62 |
| Interferon alpha | group | 5.38E-62 |
| IL10 | cytokine | 2.80E-57 |
| tretinoin | chemical - endogenous mammalian | 2.07E-56 |
| TLR4 | transmembrane receptor | 9.31E-56 |
| tetradecanoylphorbol acetate | chemical drug | 4.18E-55 |
| SB203580 | chemical - kinase inhibitor | 1.54E-48 |
| LDL | complex | 1.04E-47 |
| TLR3 | transmembrane receptor | 1.24E-46 |
| IRF7 | transcription regulator | 2.19E-46 |
| IFNB1 | cytokine | 4.67E-46 |
| IL2 | cytokine | 1.10E-45 |
| CSF2 | cytokine | 1.63E-45 |
| Immunoglobulin | complex | 2.65E-45 |
| RELA | transcription regulator | 8.58E-45 |
| TCR | complex | 1.26E-44 |
| beta-estradiol | chemical - endogenous mammalian | 1.87E-44 |
| IFNA2 | cytokine | 2.07E-44 |
| MYD88 | other | 5.17E-44 |
| OSM | cytokine | 6.37E-43 |
| E. coli B5 lipopolysaccharide | chemical - endogenous non-mammalian | 4.64E-42 |
| CD40LG | cytokine | 5.96E-42 |

**Figure 4 – figure supplement 1. Module – trait relationships in axotomized DRGs and analysis of the turquoise module**

**A**. Heatmap of the correlation of WGCNA modules with indicated experimental conditions (sham operated mice and following nerve injury). The values in each cell are Pearson's correlation co-efficient and Student asymptotic p-values (parenthesis). The green to red color represents strong negative to positive-correlation of experimental condition and Module Eigen (ME) gene expression. **B**. Induction of turquoise gene co-expression module in axotomized DRGs of wildtype (WT) mice following SNC at d1, d3 and d7 (ME gene expression). **C.** GO terms enriched in the turquoise module. Normalized enrichment scores (NES) are shown. **D**. Ingenuity pathway analysis (IPA) predicted upstream regulators of injury-induced immune pathways in axotomized DRGs. GSEA's core enrichment genes in immune pathways obtained from differentially expressed genes comparing d1 to sham DRGs; FDR<0.1 were used as input for IPA.

**Figure 5 – source data 1. List of top 100 cluster enriched genes for all cell clusters identified in the 3d post-SNC nerve.**

First column (P_val), probability of getting the "elevated" expression values these cells have under the null hypothesis that all cells have the same expression of the gene. Second column (avg_logFC), average log2 Fold-Change between cells in this cluster relative to cells in all other clusters. Third column (pct.1), percent of the cluster's cells which express the gene. Fourth column (pct.2), percent of non-cluster cells which express the gene. Fifths column (p_val_adj), p_val adjusted so that 5% of the list is expected to false positives. Links to STRING REACTOME pathway analysis for each cell cluster are included on the last page.

Figure 5 – Figure Supplement 1

A

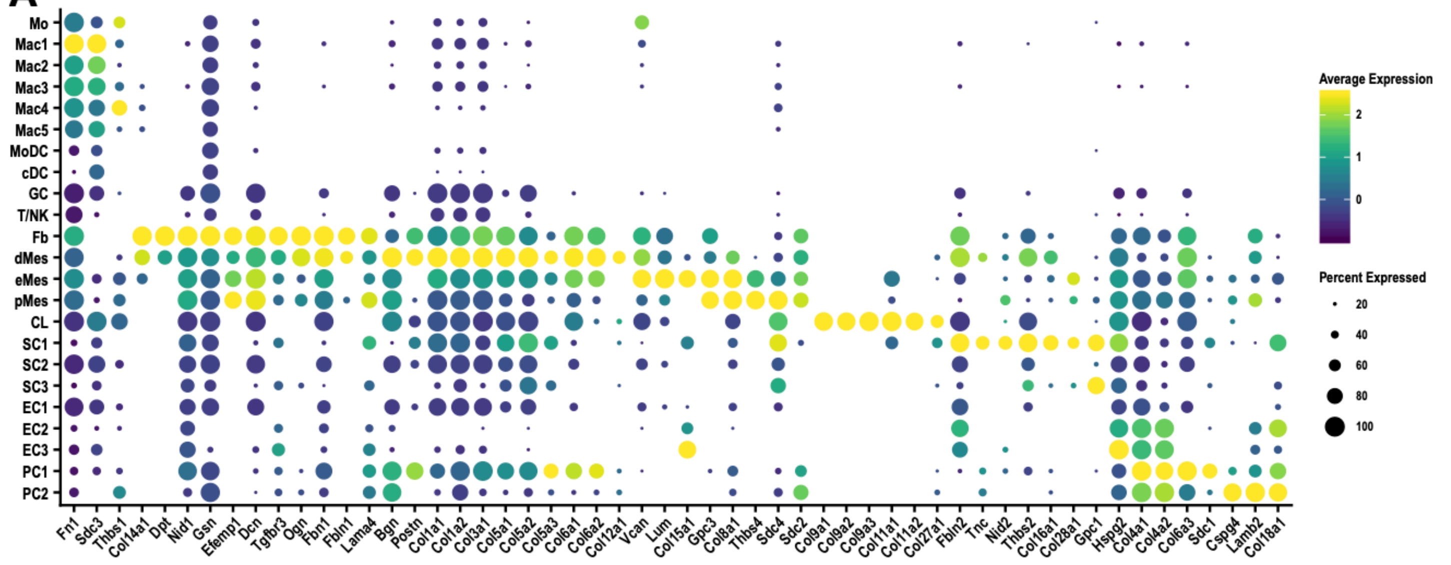


B

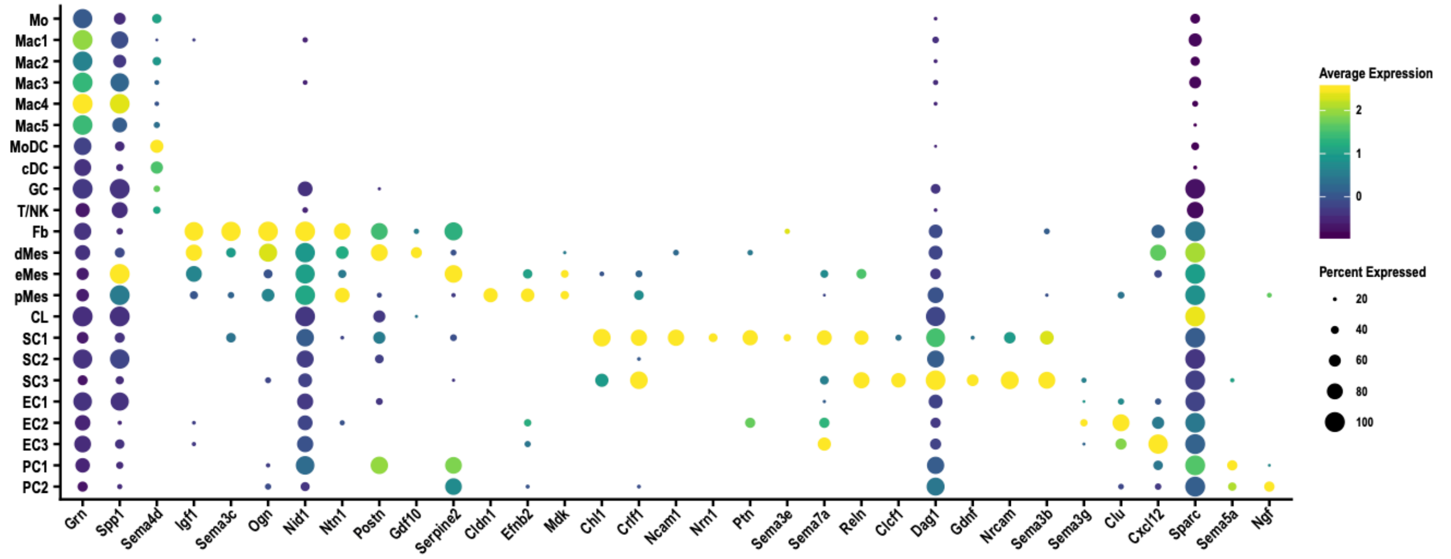

**Figure 5 – figure supplement 1. Cell cluster specific expression of ECM components and molecules that regulate axon growth in the injured sciatic nerve.**

**A**. Dotplot analysis of extracellular matrix molecules prominently expressed in the d3 post-SNC nerve. **B**. Dotplot analysis of gene products implicated in axon growth, guidance, and regeneration in the d3 post-SNC nerve. Expression levels, normalized to average gene expression (color coded) are shown. For each cell cluster the percentile of cells expressing a specific gene is indicated by the dot size.

### Figure 5 – Figure Supplement 2

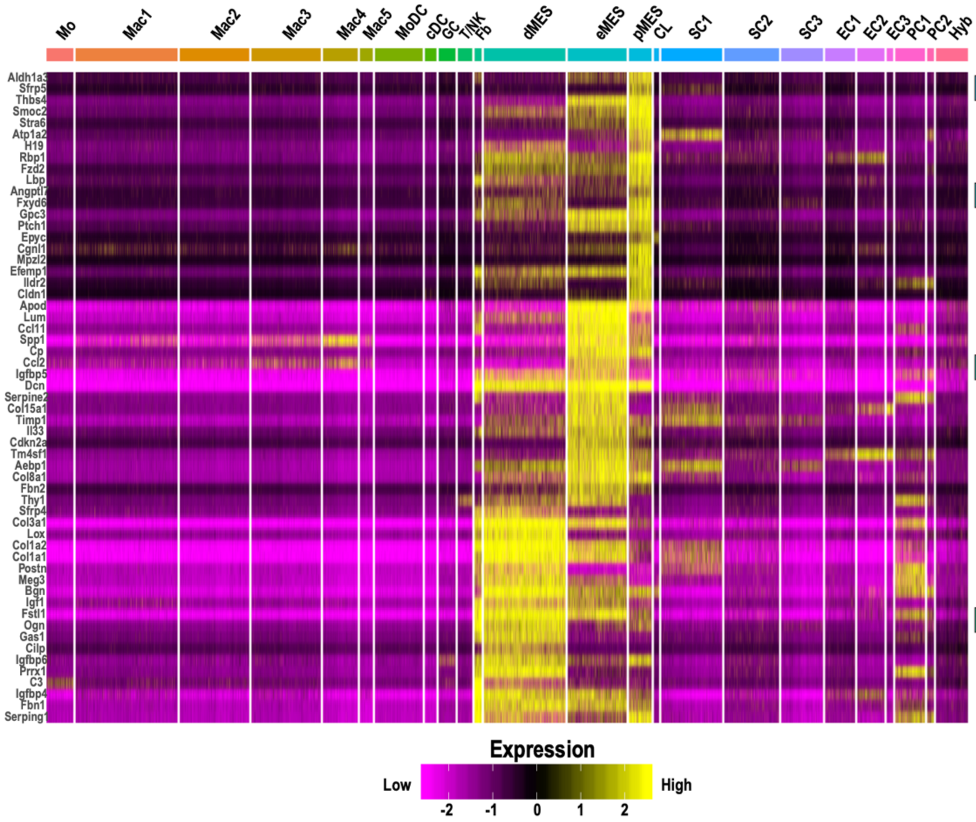


| STRING REACTOME Pathways: | FDR: |
|---|---|
| **pMES** | |
| Extracellular Matrix Organization | 5.37e-05 |
| Signaling by Receptor Tyrosine Kinases | 0.00010 |
| **eMES** | |
| Extracellular Matrix Organization | 4.21e-24 |
| Assembly of Collagen Fibrils and other Multimeric Structures | 2.68e-11 |
| Collagen Biosynthesis and Modifying Enzymes | 2.68e-11 |
| Collagen Degradation | 2.68e-11 |
| **dMES** | |
| Extracellular Matrix Organization | 8.59e-30 |
| Assembly of Collagen Fibrils and other Multimeric Structures | 2.52e-19 |
| Collagen Biosynthesis and Modifying Enzymes | 2.52e-19 |
| ECM Proteoglycans | 4.24e-18 |
| Integrin Cell Surface Interactions | 1.11e-12 |
| Regulation of Insulin-like Growth Factor (IGF) Transport and Uptake by Insulin-like Growth Factor Binding Proteins (IGFBPs) | 1.21e-12 |
| NCAM1 Interactions | 1.38e-10 |
| Axon Guidance | 2.04e-07 |
| **Fb** | |
| Extracellular Matrix Organization | 4.94e-13 |
| Degradation of the Extracellular Matrix | 6.27e-06 |
| Molecules Associated with Elastic Fibres | 2.63e-05 |
| Regulation of Complement Cascade | 5.27e-05 |

**Figure 5 – figure supplement 2. Single cell gene expression in mesenchymal cell clusters in injured sciatic nerve.**

Heatmap of top genes enriched in perineural mesenchymal cells (pMES), endoneurial mesenchymal cells (eMES), differentiating mesenchymal cells (dMES), and epineural fibroblasts (Fb) in the d3 post-SNC nerve. Expression levels are calibrated to median gene expression. STRING REACTOME Pathways are listed. FDRs (false discovery rates) are shown.

Figure 5 – Figure Supplement 3

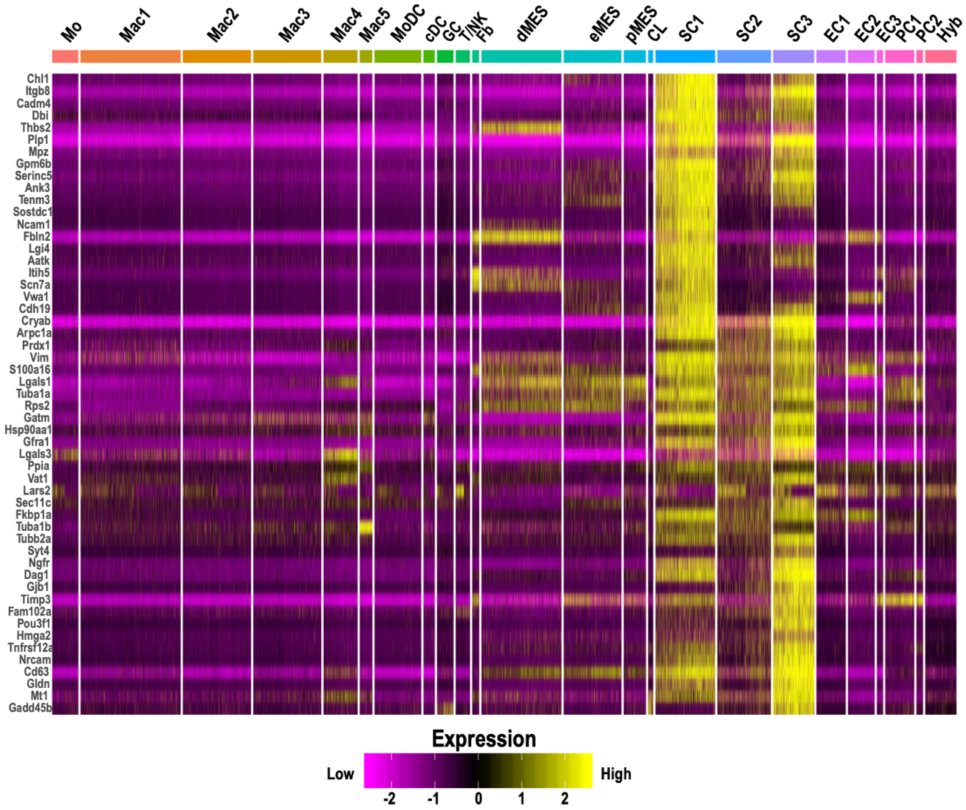


| STRING REACTOME Pathways: | FDR: |
|---|---|
| **SC1** | |
| Extracellular Matrix Organization | 2.91e-09 |
| Axon Guidance | 3.24e-05 |
| Integrin Cell Surface Interactions | 8.52e-05 |
| Collagen Formation | 0.00028 |
| **SC2** | |
| Cellular Response to Stress | 1.14e-06 |
| The Role of GTSE1 in G2/M Progression after G2 Checkpoint | 1.66e-06 |
| HSP90 Chaperone Cycle for Steroid Hormone Receptors (SHR) | 1.94e-06 |
| Microtubule-dependent Trafficking of Connexons from Golgi to the Plasma Membrane | 1.86e-05 |
| Cell Cycle | 7.09e-05 |
| **SC3** | |
| Axon Guidance | 5.30e-07 |
| Cellular Response to Stress | 2.97e-05 |
| GAP Junction Assembly | 2.97e-05 |
| L1CAM Interactions | 3.58e-05 |

**Figure 5 – figure supplement 3. Single cell gene expression in Schwann cell clusters in injured sciatic nerve.**

Heatmap of top genes enriched in repair Schwann cell clusters (SC1, SC2, and SC3) in the d3 post-SNC nerve. Expression levels are calibrated to median gene expression. STRING REACTOME Pathways are listed. FDRs (false discovery rates) are shown.

Figure 5 – Figure Supplement 4

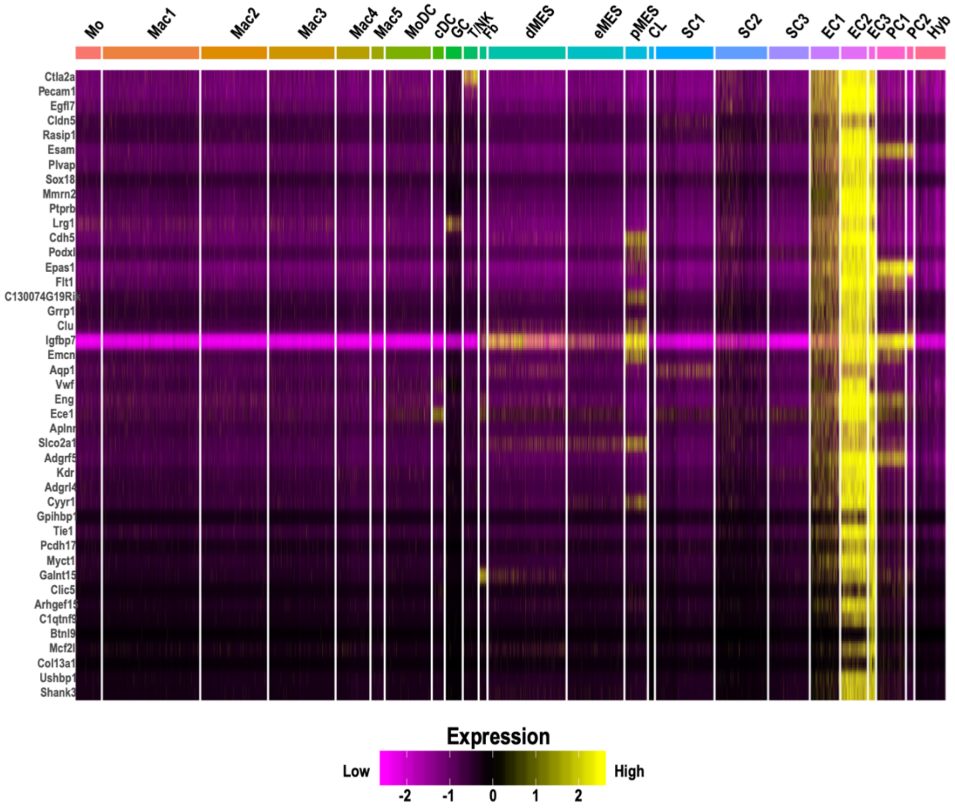


| STRING KEGG Pathways: | FDR: |
|---|---|
| **EC1** | |
| Focal Adhesion | 1.71e-13 |
| Vascular Smooth Muscle Contraction | 2.91e-08 |
| ECM-Receptor Interaction | 2.30e-07 |
| **EC2** | |
| Cell Adhesion Molecules (CAMs) | 9.24e-05 |
| Leukocyte Transendothelial Migration | 0.00098 |
| **EC3** | |
| Fluid Shear Stress and Atherosclerosis | 0.00047 |
| Leukocyte Transendothelial Migration | 0.00092 |

**Figure 5 – figure supplement 4. Single cell gene expression in endothelial cell clusters in injured sciatic nerve.**

Heatmap of top genes enriched in endothelial cell clusters (EC1, EC2, and EC3) in the d3 post-SNC nerve. Expression levels are calibrated to median gene expression. STRING KEGG Pathway analysis and FDRs (false discovery rates) are shown.

Figure 5 – Figure Supplement 5

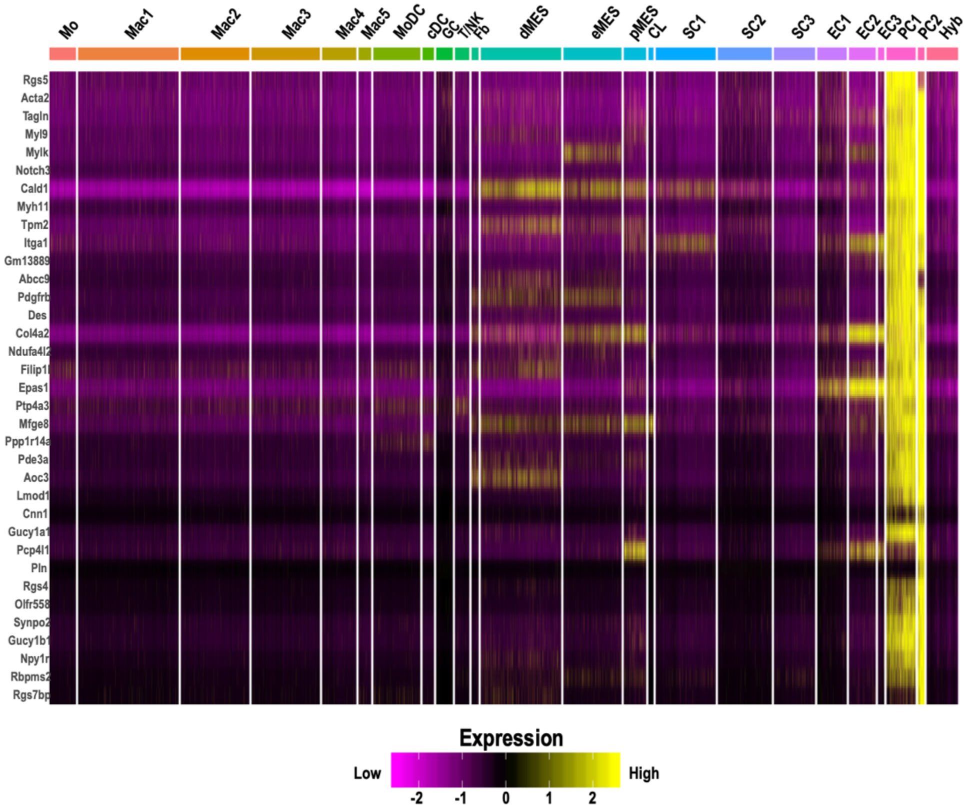


| STRING KEGG Pathways: | FDR: |
|---|---|
| **PC1** | |
| Focal Adhesion | 3.53e-12 |
| Vascular Smooth Muscle Contraction | 1.44e-09 |
| cGMP-PKG Signaling Pathway | 3.95e-05 |
| **PC2** | |
| Focal Adhesion | 1.71e-13 |
| Vascular Smooth Muscle Contraction | 2.91e-08 |
| ECM- Receptor Interaction | 2.30e-07 |

**Figure 5 – figure supplement 5. Single cell gene expression in pericyte cell clusters in injured sciatic nerve.**

Heatmap of top genes enriched in pericyte clusters (PC1 and PC2) in the d3 post-SNC nerve. Expression levels are calibrated to median gene expression. STRING REACTOME Pathways are listed. FDRs (false discovery rates) are shown.

Figure 5 – Figure Supplement

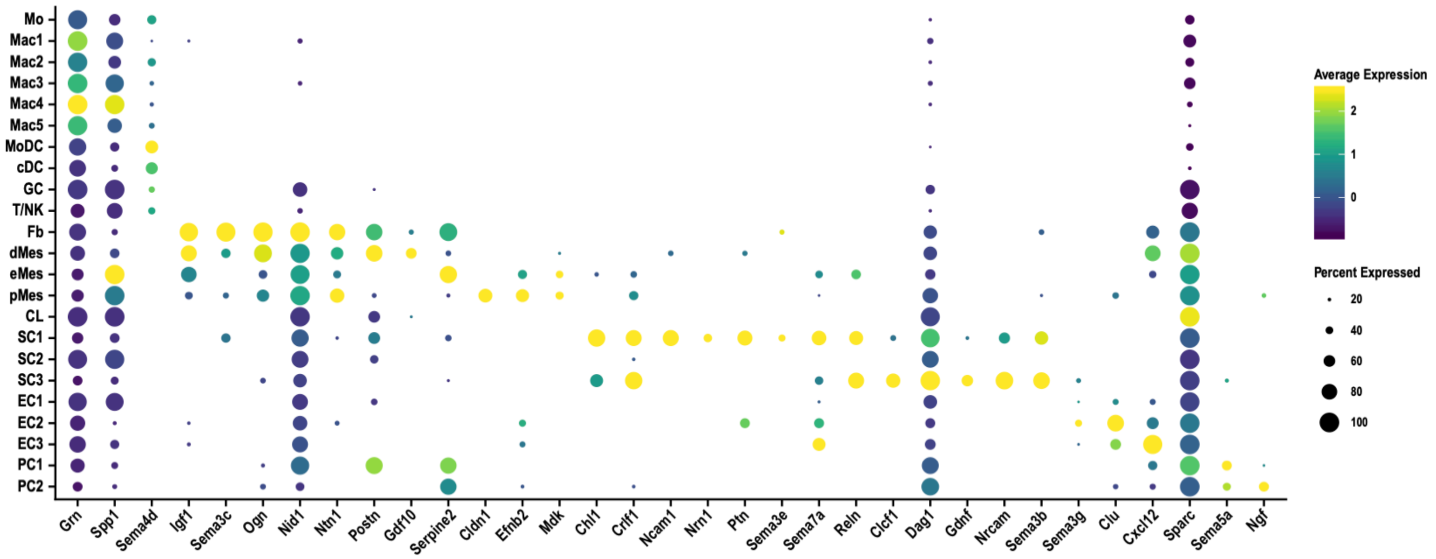

**Figure 5 – figure supplement 6. Single cell gene expression of immune modulatory molecules in the injured PNS tissue.**

Dotplot analysis of gene products with immune modulatory function in the d3 post-SNC nerve. Expression levels, normalized to average gene expression (color coded). For each cell cluster the percentile of cells that express the listed gene (dot size) is shown.

Figure 6 – Figure Supplement 1

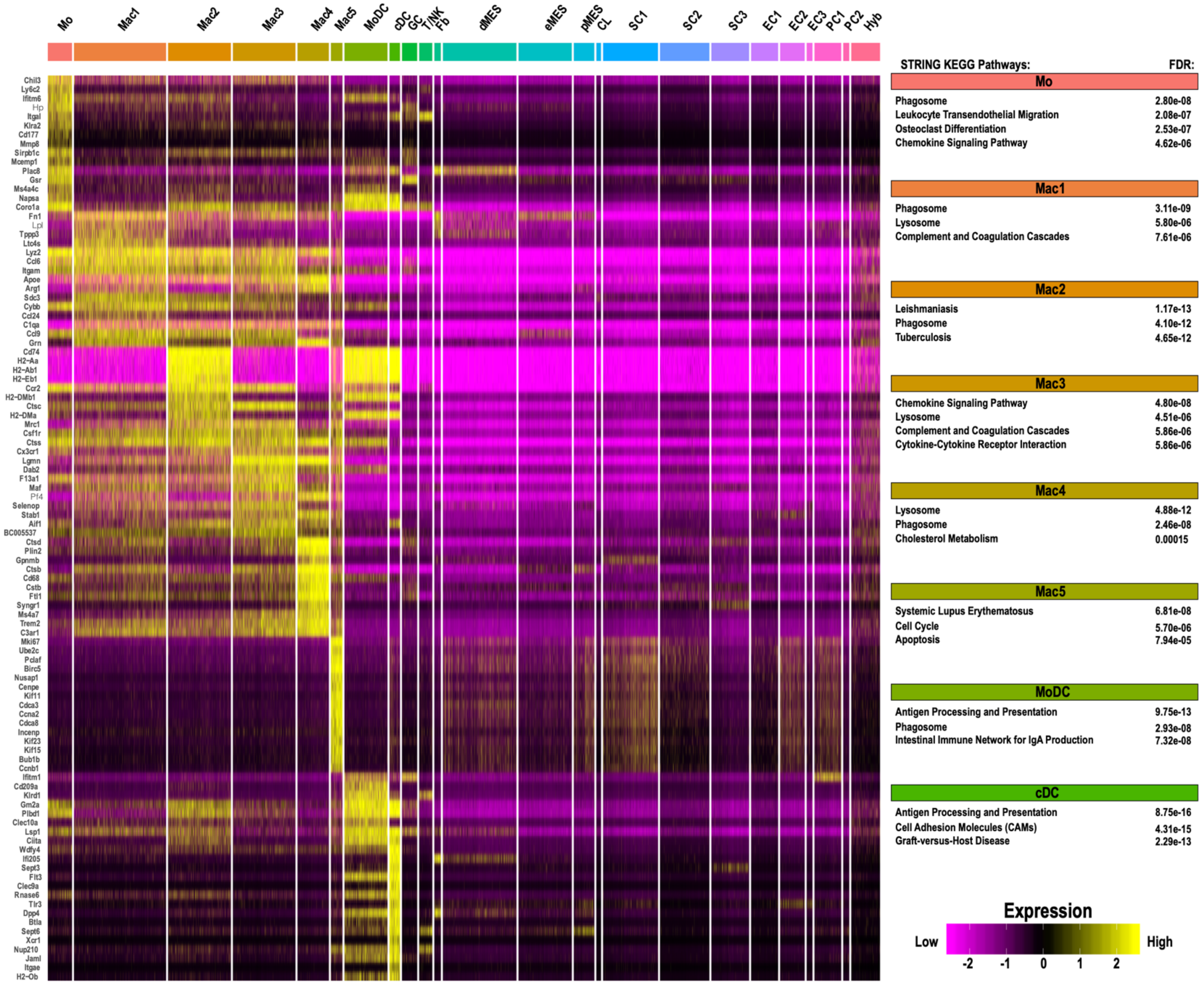

**Figure 6 – figure supplement 1. Single cell gene expression in myeloid cell clusters of injured sciatic nerve.**

Heatmap of top genes enriched in monocytes (Mo), macrophage clusters 1-5 (Mac1-5), MoDC, and cDC in the d3 post-SNC nerve. Expression levels are calibrated to median gene expression. STRING KEGG Pathway analysis and FDRs (false discovery rates) are shown.

Figure 6 – Figure Supplement 2

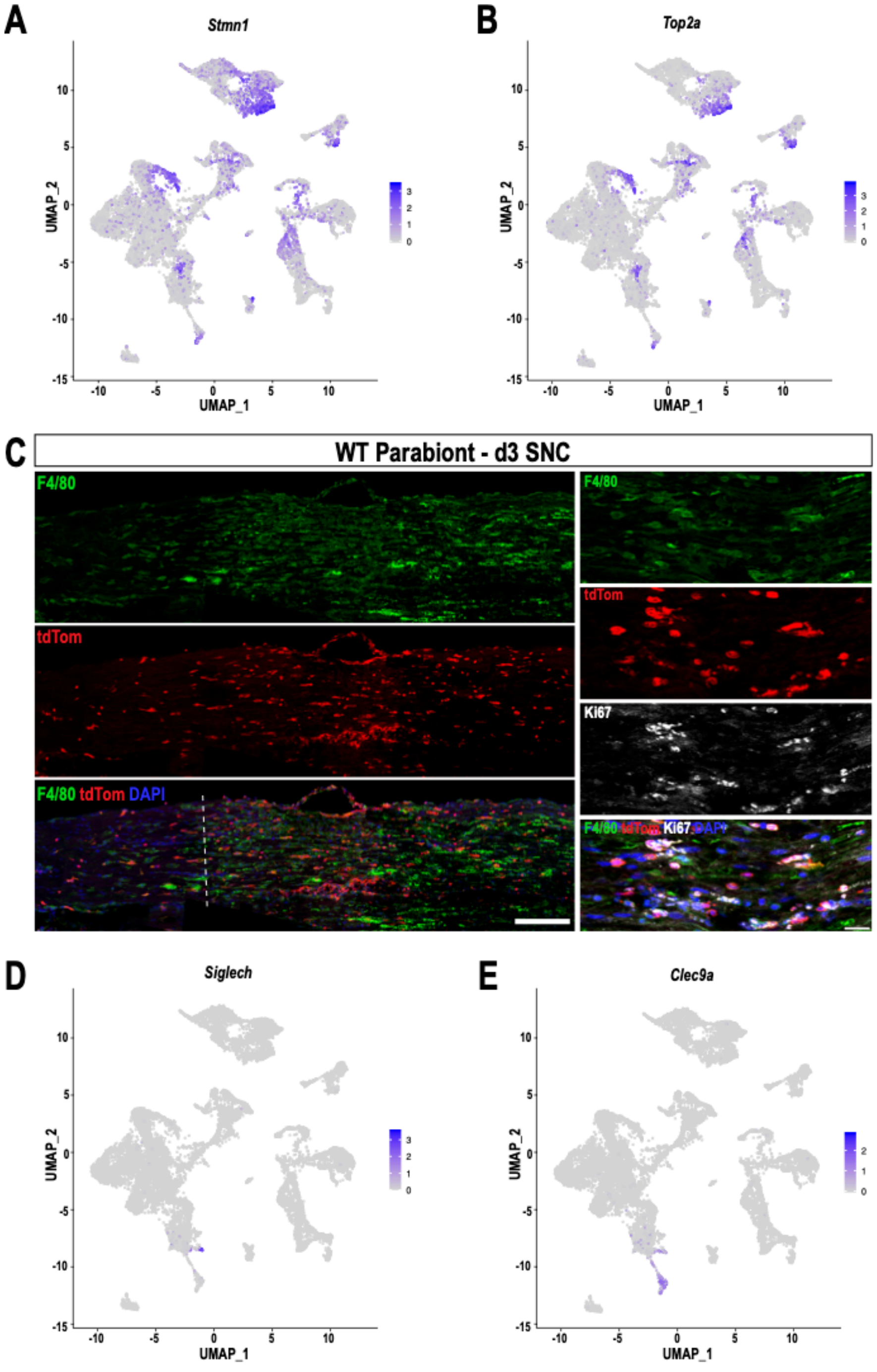

**Figure 6 – figure supplement 2. Identification of blood-borne, stem-like myeloid cells in the injured sciatic nerve**

**A**. Feature plots of *Stmn1* (stathmin-1) and **B**. *Top2a* (DNA topoisomerase II alpha) highlight proliferating cells, including Mac5 cells and a small group of myeloid cells located between clusters Mac2 and MoDC. **C**. Longitudinal sciatic nerve section of 3d WT parabiont, the dotted line marks the injury site, proximal is to the left. $TdTom^+$ cells are blood-borne immune cells originating from the tdTom parabiont. Macrophages are stained with anti-F4/80 (green), scale bar, 100 µm. Higher magnification images show the injury site. Some $tdTom^+F4/80^+$ macrophages are stained with anti-Ki67 (white), scale bar, 50 µm. **D**. Feature plots of *Siglech*, a marker for pDC and **E**. *Clec9a*, a marker for cDC, in the 3d injured nerve.

Figure 6 – Figure Supplement 3

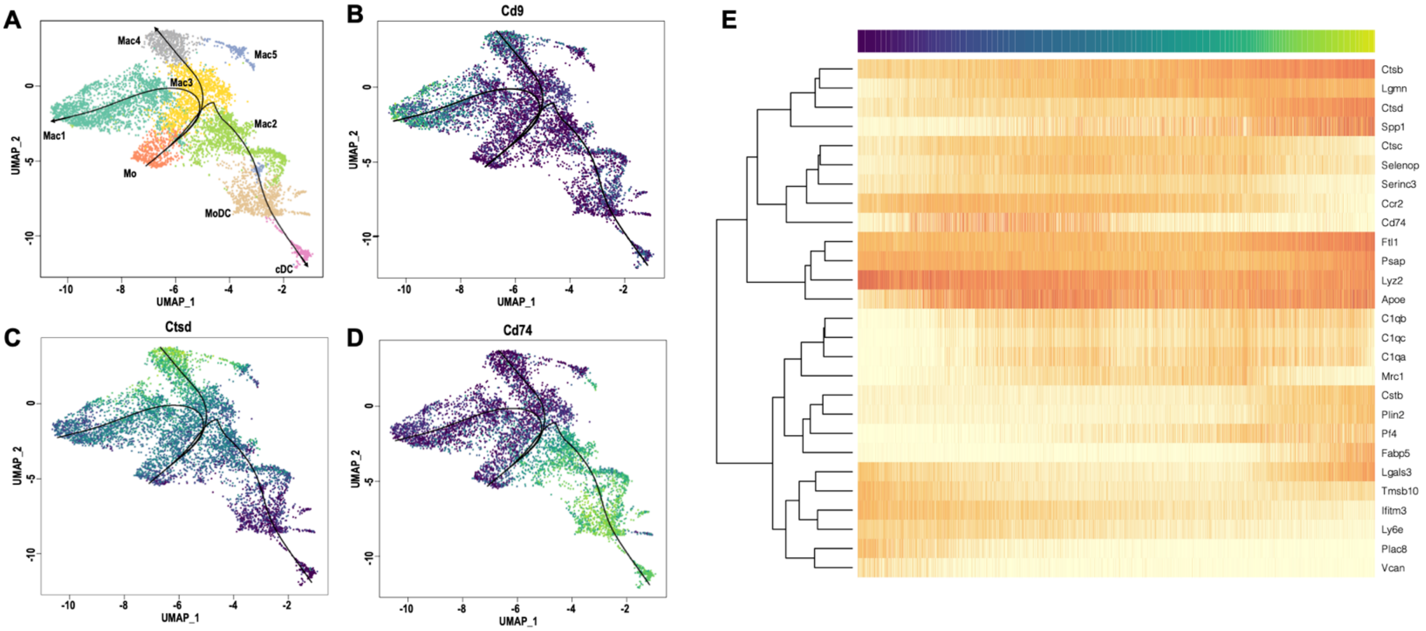

**Figure 6 – figure supplement 3. Monocyte to macrophage differentiation based on single cell expression modeling.**

Pseudo time trajectory analysis of cell differentiation in the d3 post-SNC nerve **A**. Predicted cell differentiation in the myeloid cell compartment. Slingshot analysis was used to predict how Mo (monocytes) differentiate into specific Mac subpopulations and MoDC. Feature plots for *Cd9, Ctsd,* and *Cd74* are shown as representative examples. **B**. Pseudo time gene expression changes of top genes used for trajectory analysis.

Figure 6 – Figure Supplement 4

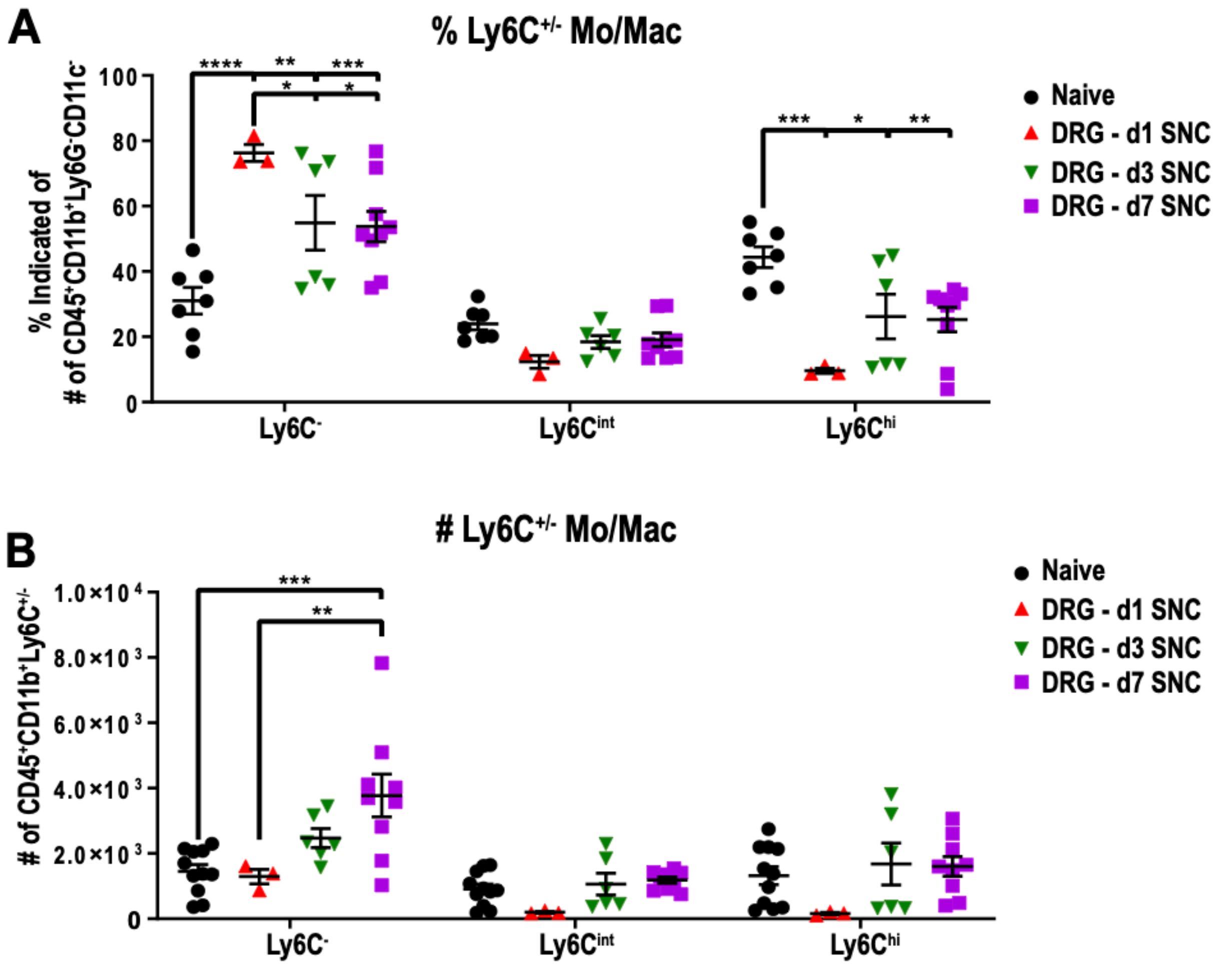

**Figure 6 – figure supplement 4. Mo/Mac maturation in axotomized DRGs, assessed by Ly6C surface expression.**

Flow cytometric analysis of Mo/Mac in naive and axotomized DRGs. The surface distribution of Ly6C on Mo/Mac ($CD45^+$, $CD11b^+$, $Ly6G^-$, $CD11c^-$) is shown. **A**. The percentile of Ly6C cells and **B**. the number of Ly6C cells in naive DRGs, d1, d3, and d7 post-SNC. Mo/Mac were binned into $Ly6C^{hi}$, $Ly6C^{int}$, and $Ly6C^-$ cells. Flow data are represented as mean ± SEM. Each data point represents L3-L5 DRGs pooled from 3-4 animals (18-24 DRGs), biological replicates, n= 3-9. Statistical analysis was performed in GraphPad Prism (v7) using 1-way or 2-way ANOVA with correction for multiple comparisons with Tukey's post-hoc test. A p value < 0.05 (*) was considered significant. p < 0.01 (**), p < 0.001 (***), and p < 0.0001 (****).

Figure 6 – Figure Supplement 5

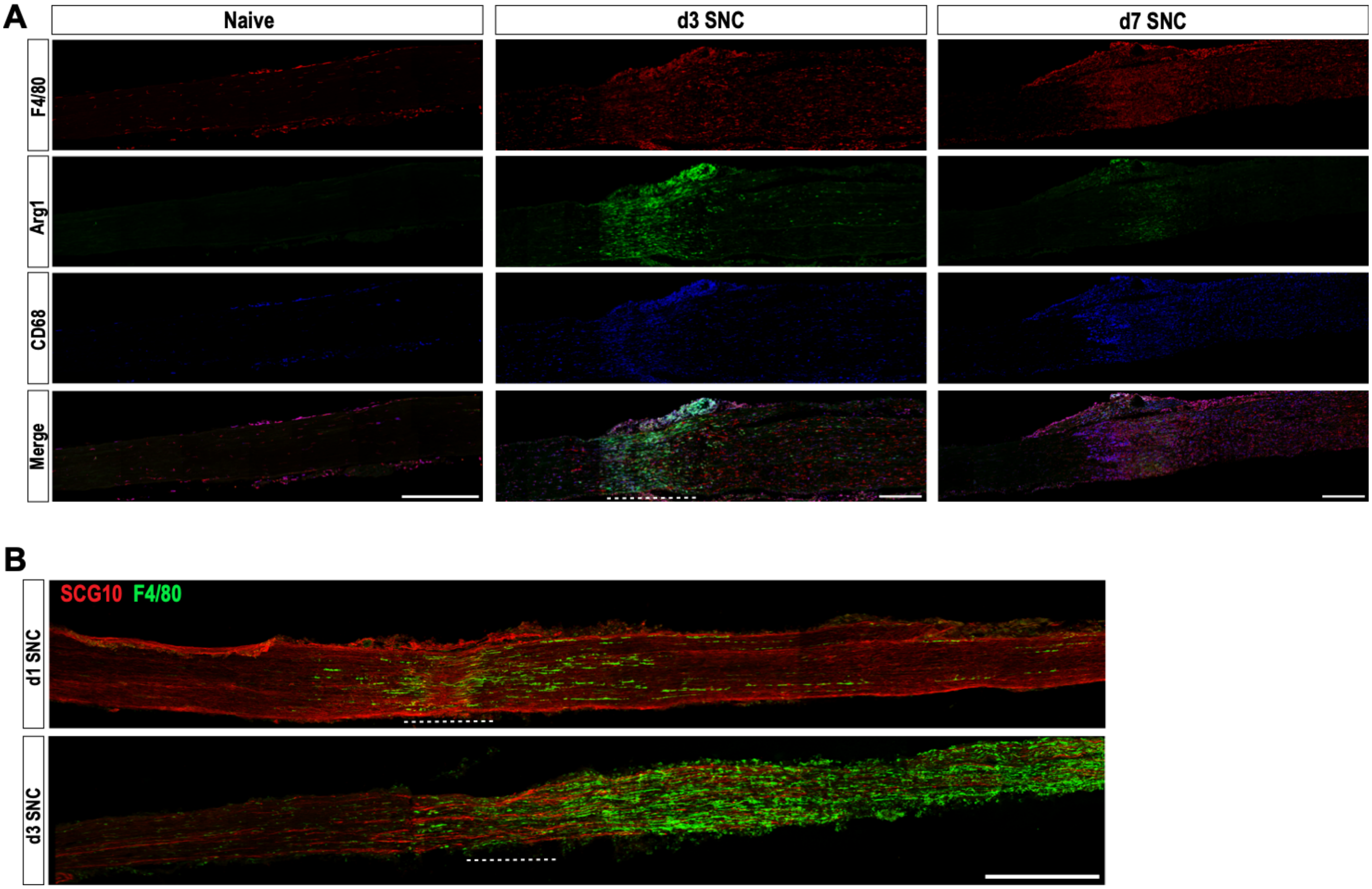

**Figure 6 – figure supplement 5. Evidence for distinct immune compartments in the injured sciatic nerve**

**A**. Longitudinal sections of naïve and injured (3d and 7d) sciatic nerves from *Arg1-YFP* reporter mice. The injury site is underlined with a dotted line. Proximal is to the left. Distribution of F4/80$^+$, *Arg1-YFP*$^+$, and CD68$^+$ macrophages is shown. Scale bar, 200 µm. **B**. Longitudinal sciatic nerve sections of adult mice at d1 and d3 post-SNC. The crush site is marked with a dotted line, proximal is to the left. Nerves were stained with anti-SCG10 to visualize regenerating sensory axons and anti-F4/80 to stain for a subpopulation of macrophages. At d1, F4/80$^+$ macrophages are found near the crush site. At d3, F4/80$^+$ macrophages are abundantly present throughout the distal nerve stump, adjacent to SCG10$^+$ regenerating sensory axons. Scale bar, 500 µm.

Figure 6 – Figure Supplement 6

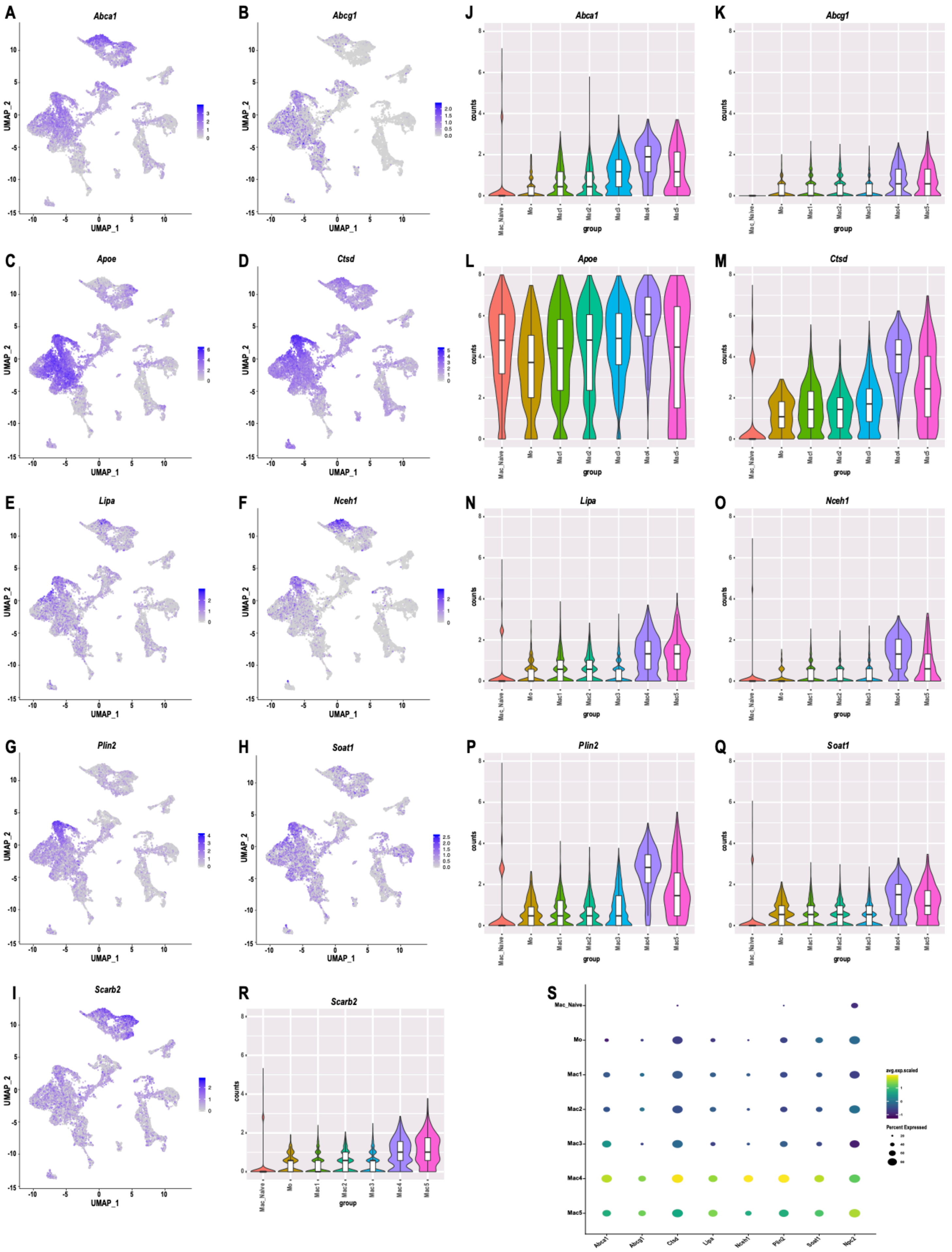

A
Abca1
B
Abcg1
C
Apoe
D
Ctsd
E
Lipa
F
Nceh1
G
Plin2
H
Soat1
I
Scarb2
J
K
L
M
N
O
P
Q
R
S
UMAP_1
UMAP_2
counts
group
Mac_Naive
Mo
Mac1
Mac2
Mac3
Mac4
Mac5
Ctsd
Lipa
Nceh1
Plin2
Soat1
Npc2
avg.exp.scaled
Percent Expressed

**Figure 6 – figure supplement 6. Expression of gene products that regulate cholesterol transport and metabolism in macrophages of injured nerve and naïve nerve**

Feature plots of gene products implicated in cholesterol transport and lowering of cellular cholesterol levels. **A.** *Abca1/CERP* (cholesterol efflux regulatory protein). **B.** *Abcg1* (ATP-binding cassette subfamily G). **C.** *Apoe* (apolipoprotein E). **D.** *Ctsd* (cathepsin D). **E.** *Lipa* (lipase A). **F.** *Nceh1* (neutral cholesterol ester hydrolase 1). **G.** *Plin2* (Perilipin 2). **H.** *Soat1* (sterol O-acyltransferase 1), **I**. *Scarb2* (scavenger receptor class B 2). Calibration of gene expression for each plot is shown. **J-R**. Violin plots for cholesterol regulatory gene products in naïve Mac, Mo and Mac1-5. **S.** Dot plot showing expression of cholesterol transporters and metabolic enzymes in naïve nerve Mac in comparison to Mo and Mac1-5 in injured nerve.  Expression levels are normalized to average gene expression (color coded). For each cell cluster the percentile of cells that express the listed gene (dot size) is shown.

Figure 7 – Figure Supplement 1

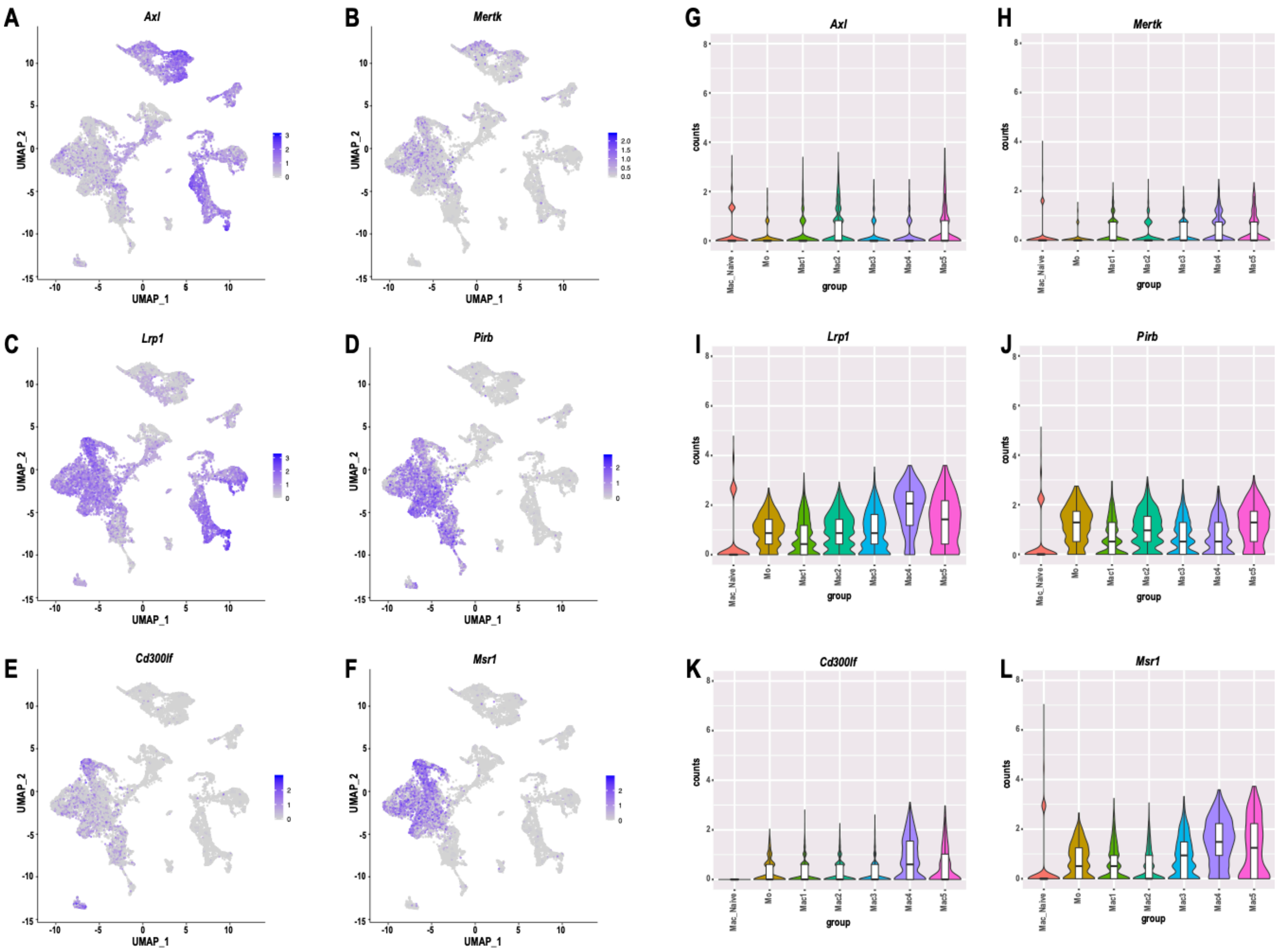

**Figure 7 – figure supplement 1. Expression of gene products implicated in myelin binding and phagocytosis**

**A-F.** Feature plots and **G-L.** Violin plots of *Axl* (TAM receptor tyrosine kinase AXL), *Mertk* (TAM receptor tyrosine kinase MER)*, Lrp1* (Low density lipoprotein receptor-related protein 1)*, Pirb* (Paired Ig-like receptor B) *Cd300lf* (CD300 Molecule Like Family Member F)*, and Msr1* (Macrophage scavenger receptor 1) expression in the d3 post-SNC nerve. Violin plots are shown for Mac from naïve nerve, Mo, and Mac1-5 from 3d post-SNC nerves.

Figure 7 – Figure Supplement 2

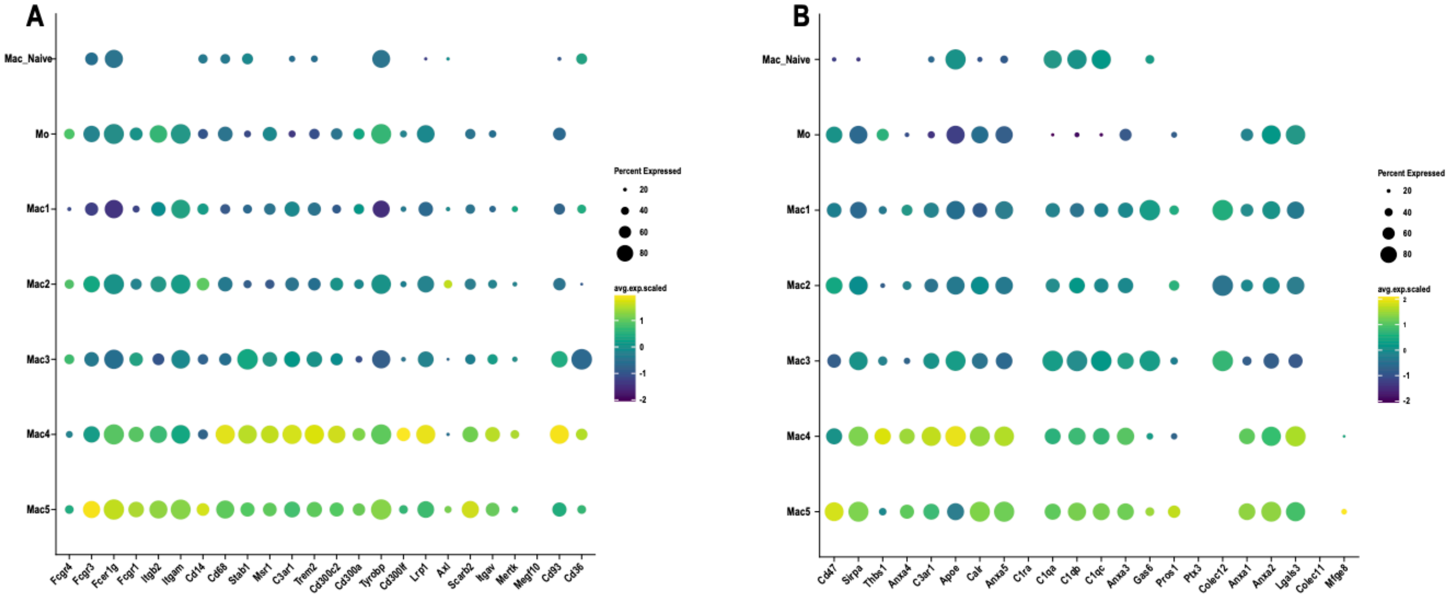

**Figure 7 – figure supplement 2. Expression analysis of bridging molecules and engulfment receptors in macrophages of naïve nerve and injured nerve**

**A.** Dotplot showing expression of bridging molecules in in naïve Mac in comparison to Mo and Mac1-5 in injured nerve. **B**. Dotplot showing expression of engulfment receptors in in naïve Mac in comparison to Mo and Mac1-5 in injured nerve. Expression levels are normalized to average gene expression (color coded). For each cell cluster the percentile of cells that express the listed gene (dot size) is shown.

Figure 7 – Figure Supplement 3

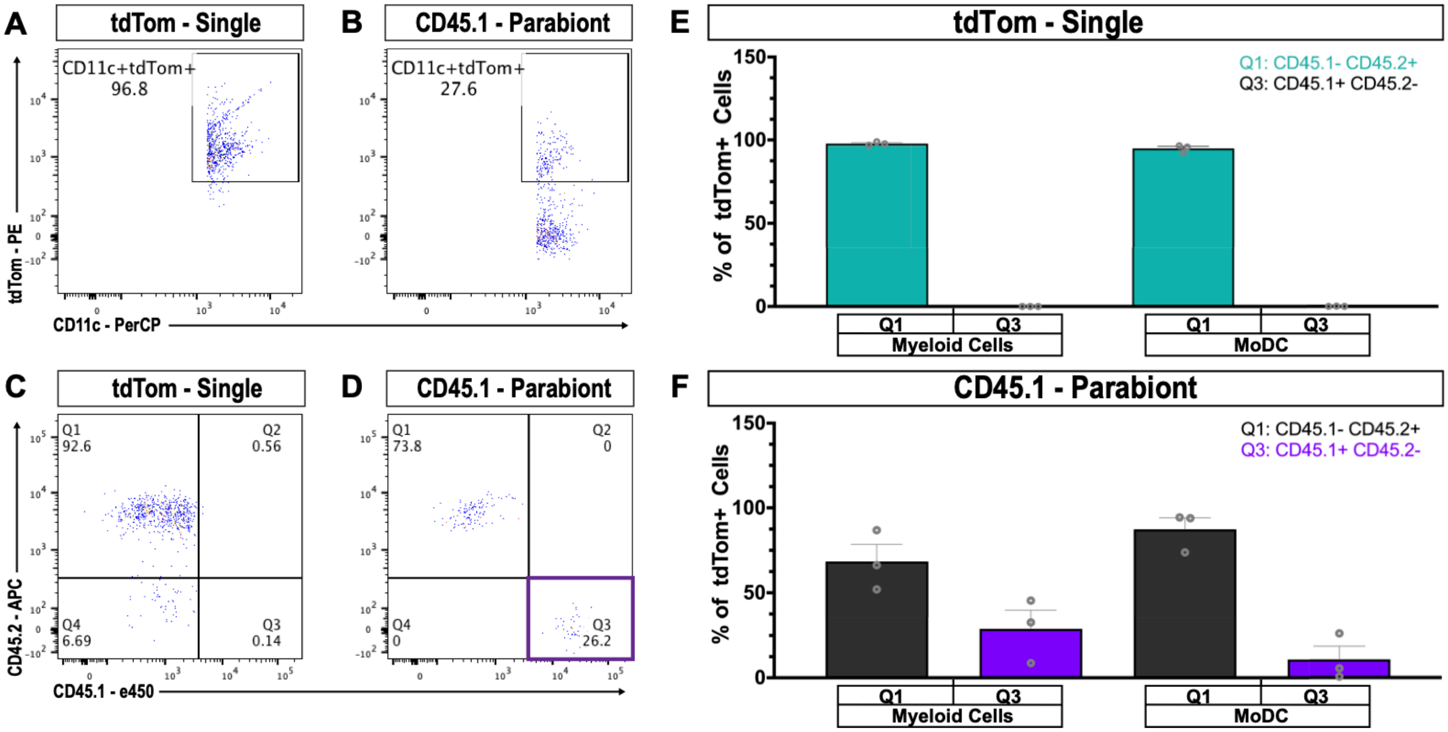

**Figure 7 – figure supplement 3. Contribution of MoDC to efferocytosis in the injured sciatic nerve**

**A.** Flow cytometric analysis of 3d sciatic nerve from non-parabiotic (single) tdTom mice. Dot plot shows that 96.8% of MoDC (CD11b$^{+}$, CD11c$^{+}$, Ly6G$^{-}$) are tdTom$^{+}$ **B.** Flow cytometric analysis of 3d sciatic nerves from CD45.1 parabionts. Dot plot shows that 27.6% of MoDC are tdTom$^{+}$ **C.** In the 3d sciatic nerves of non-parabiotic (single) tdTom mice, no tdTom$^{+}$, CD45.1$^{+}$ MoDC are present. **D**. In the 3d sciatic nerve of the CD45.1 parabiont, CD45.1$^{+}$ and CD45.2$^{+}$ MoDC are found. **E.** Quantification of CD45.1 and CD45.2 myeloid cells (CD11b$^{+}$) and MoDC in the 3d sciatic nerve of non-parabiotic (single) tdTom mice. As expected, CD45.1$^{+}$ cells are not detected. **F**. Quantification of CD45.1 and CD45.2 myeloid cells (CD11b$^{+}$) and MoDC in the 3d sciatic nerve of the CD45.1 parabiont. The presence of tdTom$^{+}$, CD45.1$^{+}$, CD45.2$^{-}$, CD11b$^{+}$, CD11c$^{+}$, Ly6G$^{-}$ cells, indicates that MoDC participate in efferocytosis.

Figure 8 – Figure Supplement 1

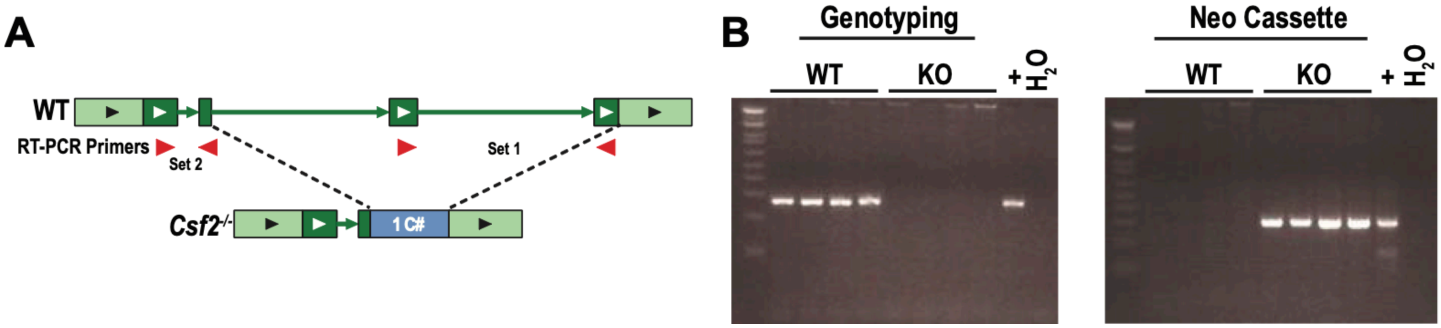

**Figure 8 – figure supplement 1. Locus and PCR genotyping of *Csf2*$^{-/-}$ mice.**

**A.** Schematic of *Csf2* gene locus of WT and germline *Csf2*$^{-/-}$ mice. Primer sets were designed both inside (set 1) and outside (set 2) of the deleted region. **B.** Agarose gel shows PCR genotyping results from naïve brains of WT and *Csf2*$^{-/-}$ (KO) animals against primer set 2 (left) or against the neomycin cassette (right). In the absence of genomic DNA (water ($H_2O$) control), no PCR product is observed.
